# Subregion-Specific Input Organization of Prefrontal-Projecting Basal Forebrain Cholinergic Neurons and Weakened Striatal–NBM Inhibitory Transmission in 5xFAD mice

**DOI:** 10.64898/2026.06.11.731708

**Authors:** Yufei Huang, Xueyi Xie, Michael Fernaine, Ziyi Li, Xuehua Wang, Jun Wang

## Abstract

Basal forebrain cholinergic neurons regulate cortical activity and cognition and are vulnerable in Alzheimer’s disease (AD). However, the upstream circuits controlling projection-defined basal forebrain cholinergic populations remain incompletely understood. Here, we used projection-specific rabies-mediated monosynaptic tracing to map whole-brain inputs to medial prefrontal cortex (mPFC)-projecting cholinergic neurons in the nucleus basalis of Meynert (NBM) and horizontal limb of the diagonal band of Broca (HDB). mPFC-projecting NBM and HDB cholinergic neurons received broad but distinct input patterns. NBM cholinergic neurons received prominent striatal input, including input from D1-expressing medium spiny neurons, whereas HDB cholinergic neurons showed proportionally weaker striatal input and broader non-striatal contributions. Optogenetic electrophysiology confirmed that striatal inputs formed monosynaptic GABAergic inhibitory synapses onto NBM cholinergic neurons. This inhibitory transmission was weakened in 5xFAD mice, indicating impairment of a striatal–NBM inhibitory circuit in an AD mouse model. Together, these findings reveal subregion-specific input organization of mPFC-projecting basal forebrain cholinergic neurons and identify a vulnerable striatal–NBM circuit in AD.

**Highlights:**

1. Whole-brain rabies tracing reveals input organization of mPFC-projecting BF cholinergic neurons.
2. NBM and HDB cholinergic neurons projecting to mPFC show distinct monosynaptic input profiles.
3. Striatal D1-MSNs are a major input source to mPFC-projecting NBM cholinergic neurons.
4. Striatal–NBM inhibitory transmission is functionally impaired in 5xFAD mice.

## INTRODUCTION

The basal forebrain (BF) is a major neuromodulatory system that regulates cortical activity through widespread cholinergic projections (Mesulam et al., 1983; Sarter et al., 2003; Hasselmo and Sarter, 2011; Ballinger et al., 2016; Ananth et al., 2023). BF cholinergic neurons play critical roles in attention, learning and memory, arousal, sensory processing, and cognitive flexibility by modulating cortical and hippocampal circuits. Dysfunction of BF cholinergic signaling has been implicated in several neurological and psychiatric disorders, including Alzheimer’s disease (AD), schizophrenia, and addiction (Hyde and Crook, 2001; Williams and Adinoff, 2008; Avram et al., 2021). In AD, degeneration and dysfunction of BF cholinergic neurons, particularly those in the nucleus basalis of Meynert (NBM), are closely associated with cortical cholinergic deficits and cognitive impairment (Whitehouse et al., 1982; Schmitz et al., 2018). Thus, understanding how BF cholinergic neurons are regulated by their upstream inputs is essential for defining the circuit mechanisms that shape cortical function in health and disease.

The BF is anatomically and neurochemically heterogeneous. Basal forebrain cholinergic neurons are distributed across multiple subregions, including the medial septum, vertical and horizontal limbs of the diagonal band of Broca, substantia innominata, and nucleus basalis of Meynert (Mesulam et al., 1983; Chaves-Coira et al., 2018; Liu et al., 2018). These subregions contain partially overlapping but distinct cholinergic projection populations that innervate different cortical and limbic targets. The NBM provides prominent cholinergic innervation to broad neocortical regions and has been strongly associated with cortical activation, pain, attentional processing, and cognitive function (Price and Stern, 1983; Zaborszky et al., 2015; Oswald et al., 2022). The HDB also contributes to cortical and limbic cholinergic innervation, including projections to prefrontal and olfactory-related regions (Ghashghaei and Barbas, 2001; Bloem et al., 2014b). This anatomical organization suggests that cholinergic neurons in different BF subregions may receive distinct afferent inputs and transmit different information streams to shared or overlapping cortical targets. However, the input organization of projection-defined BF cholinergic neurons remains incompletely understood.

The medial prefrontal cortex (mPFC) is a major target of BF cholinergic innervation and is critically involved in executive control, decision-making, attention, motivation, and emotional regulation (Miller and Cohen, 2001; Euston et al., 2012). Cholinergic modulation of the mPFC can influence neuronal excitability, cortical oscillations, and behavioral performance (Pafundo et al., 2013; Bloem et al., 2014a). Previous anatomical and physiological studies have shown that BF cholinergic neurons are influenced by multiple cortical and subcortical regions, including cortex, striatum, thalamus, hypothalamus, amygdala, and brainstem nuclei (Do et al., 2016; Hu et al., 2016; Gielow and Zaborszky, 2017). However, it remains unclear whether NBM and HDB cholinergic neurons projecting to the same cortical target, the mPFC, receive similar or distinct monosynaptic inputs.

Genetically targeted monosynaptic rabies tracing provides a powerful approach for mapping direct presynaptic inputs to defined neuronal populations (Reardon et al., 2016). By combining Cre-dependent helper-virus expression with EnvA-pseudotyped, glycoprotein-deleted rabies virus, this approach allows monosynaptic input mapping from genetically specified starter cells (Wickersham et al., 2007; Wall et al., 2010; Callaway and Luo, 2015). In the present study, we used projection-specific rabies tracing to map whole-brain monosynaptic inputs to mPFC-projecting cholinergic neurons in the NBM and HDB. We found that these two basal forebrain cholinergic populations receive broad but distinct input patterns, with a prominent striatal input to mPFC-projecting NBM cholinergic neurons. Guided by this anatomical finding, we further used optogenetic electrophysiology to examine the functional striatal–NBM connection and found that striatal inhibitory transmission onto NBM cholinergic neurons is weakened in 5xFAD mice. Our findings suggest subregion-specific input organization of mPFC-projecting basal forebrain cholinergic neurons and identify a striatal–NBM inhibitory pathway that is vulnerable in an Alzheimer’s disease mouse model.

## RESULTS

### Rabies virus-mediated retrograde monosynaptic tracing of NBM cholinergic neurons projecting to the mPFC

To identify monosynaptic inputs to nucleus basalis of Meynert (NBM) cholinergic neurons projecting to the medial prefrontal cortex (mPFC), we used a projection-specific rabies tracing strategy in ChAT-Cre;D1-tdTomato mice (Figure 1A). A 1:1 mixture of AAV-DIO-TVA-mCherry and AAV-DIO-oG was injected into the NBM to express TVA-mCherry and optimized rabies glycoprotein (oG) in Cre-expressing neurons. Twenty-one days later, EnvA-ΔG-rabies-GFP (Reardon et al., 2016) (Rabies-GFP) was injected into the mPFC. This approach allowed Rabies-GFP to infect TVA-expressing axon terminals of NBM cholinergic neurons projecting to the mPFC, retrogradely label their somata in the NBM, and spread transsynaptically to directly connected presynaptic neurons in an oG-dependent manner.

**Figure 1.**
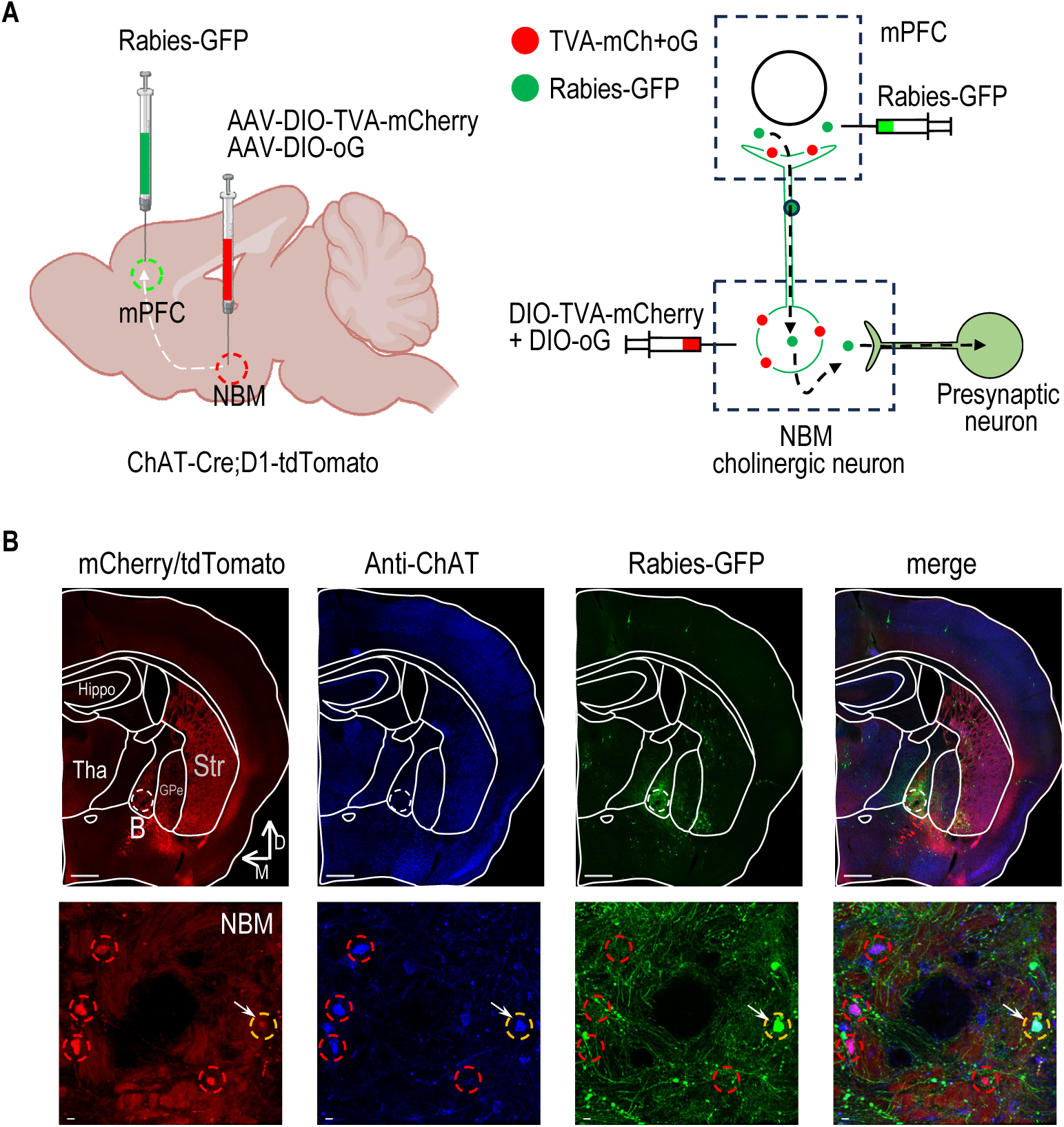
Rabies-mediated retrograde monosynaptic tracing of medial prefrontal cortex-projecting nucleus basalis of Meynert cholinergic neurons. A, Schematic showing the viral injection strategy and tracing principle. Left, a 1:1 mixture of AAV-DIO-TVA-mCherry and AAV-DIO-oG was injected into the nucleus basalis of Meynert (NBM) of ChAT-Cre;D1-tdTomato mice to selectively express TVA-mCherry and optimized rabies glycoprotein (oG) in cholinergic neurons. Twenty-one days later, EnvA-ΔG-rabies-GFP (Rabies-GFP) was injected into the medial prefrontal cortex (mPFC), the projection target of NBM cholinergic neurons. Right, Rabies-GFP selectively infected TVA-expressing axon terminals of NBM cholinergic neurons in the mPFC, was retrogradely transported to their somata in the NBM, and spread transsynaptically to directly connected presynaptic neurons in an oG-dependent manner. B, Representative images showing helper-virus expression, ChAT immunostaining, Rabies-GFP labeling, and merged fluorescence in the NBM. Low-magnification images show the injection region in the basal forebrain and Rabies-GFP-labeled cells. High-magnification images confirm the presence of starter cells. Red dashed circles indicate cholinergic neurons expressing TVA-mCherry/oG, and yellow dashed circles indicate starter cells co-expressing TVA-mCherry/oG and Rabies-GFP. Scale bars, 500 μm for low-magnification images and 10 μm for high-magnification images.

We next used ChAT immunostaining to independently verify the cholinergic identity of helper-virus-targeted neurons and Rabies-GFP-labeled starter cells in the NBM (Figure 1B). Because helper-virus-derived mCherry and endogenous D1-tdTomato fluorescence were detected in the same red channel, ChAT staining provided an independent cholinergic marker to distinguish cholinergic starter cells from nearby D1-tdTomato-positive neurons. ChAT immunostaining revealed a dense population of cholinergic neurons within the NBM. A subset of ChAT-positive neurons overlapped with red fluorescence, confirming helper-virus targeting within the NBM cholinergic population. Among these ChAT-positive, red-labeled neurons, a subset was also positive for Rabies-GFP and was therefore identified as starter cells. In addition, GFP-positive neurons lacking both ChAT immunostaining and red fluorescence were observed locally, consistent with local presynaptic neurons labeled by transsynaptic rabies spread from NBM cholinergic starter cells.

Together, these results verified the projection-specific targeting of mPFC-projecting NBM cholinergic neurons and established the tracing strategy for subsequent whole-brain mapping of their monosynaptic inputs.

### Whole-brain mapping of monosynaptic inputs to NBM cholinergic neurons projecting to the mPFC

After validating starter-cell labeling in the NBM, we examined the whole-brain distribution of Rabies-GFP-labeled presynaptic neurons to identify direct inputs to mPFC-projecting NBM cholinergic neurons. Representative coronal sections arranged from rostral to caudal revealed Rabies-GFP-labeled neurons across multiple brain regions (Figure 2A). GFP-labeled neurons were observed in the major regions quantified in this study, including the cortex, striatum, septum, thalamus, hypothalamus, hippocampus, and amygdala. Additional labeled neurons in regions not analyzed as separate categories were pooled into the “Others” group for quantification.

**Figure 2.**
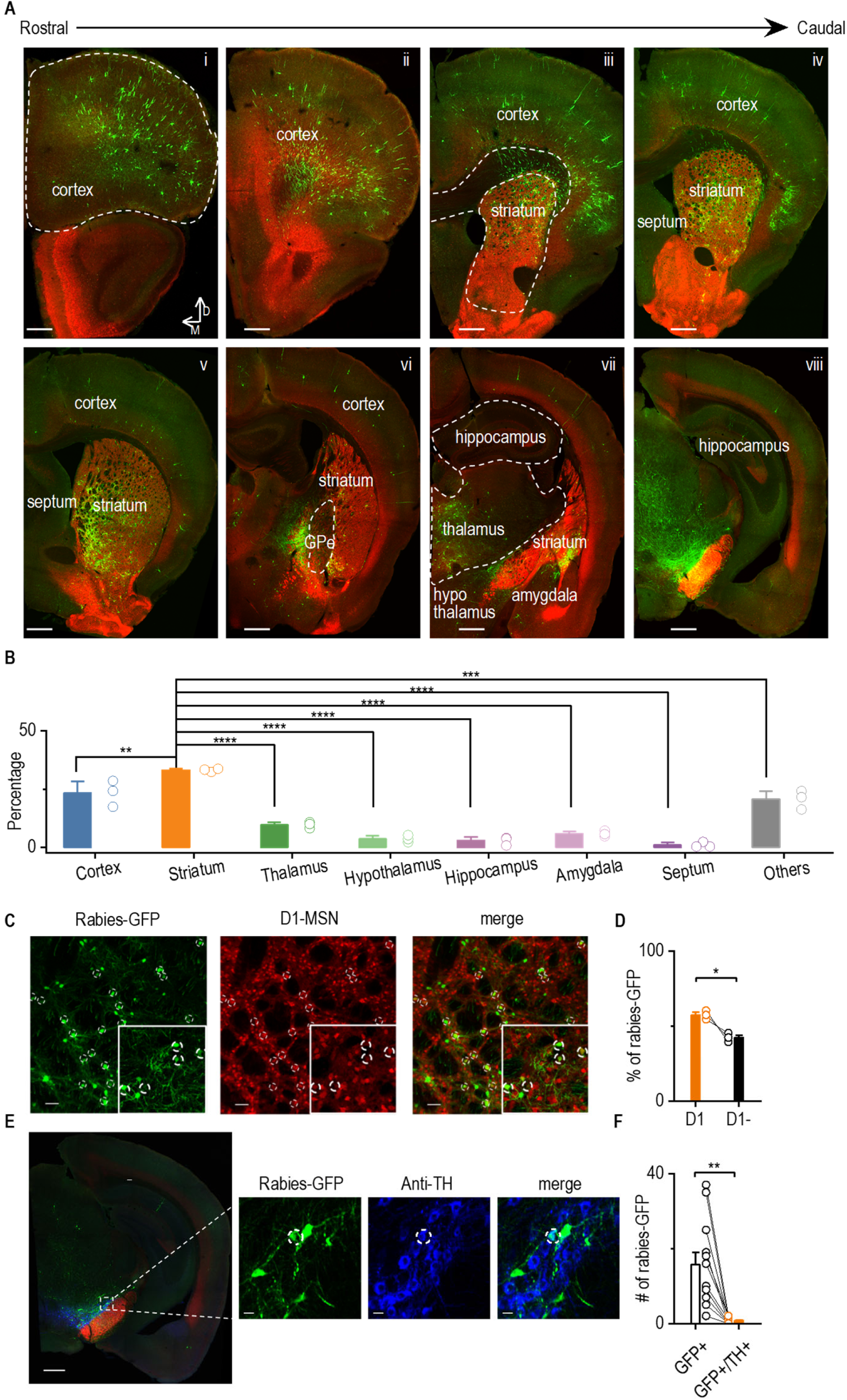
Whole-brain mapping of monosynaptic inputs to mPFC-projecting NBM cholinergic neurons. **A,** Representative coronal sections arranged from rostral to caudal showing Rabies-GFP-labeled monosynaptic input neurons throughout the brain. Cortex in sections i–vi, striatum in sections iii–vii, septum in sections iv–v, thalamus in section vii, hypothalamus in section vii, hippocampus in sections vii–viii, and amygdala in section vii. Additional labeled regions not analyzed as separate categories were pooled into the “Others” group for quantification. Dashed outlines indicate selected anatomical boundaries. Scale bars: 500 μm. B, Quantification of the relative distribution of Rabies-GFP-labeled input neurons across major brain regions. The striatum accounted for the largest proportion of monosynaptic inputs to mPFC-projecting NBM cholinergic neurons, followed by the cortex. Regions not individually quantified were included in the “Others” category. One-way ANOVA, *F*_(7, 14)_ = 50.32, *p* < 0.0001; Tukey’s post hoc comparisons involving the striatum showed significant differences between cortex and striatum (q = 6.276, *p* = 0.0076), striatum and thalamus (q = 15.16, *p* < 0.0001), striatum and hypothalamus (q = 19.05, *p* < 0.0001), striatum and hippocampus (q = 19.53, *p* < 0.0001), striatum and amygdala (q = 17.59, *p* < 0.0001), striatum and septum (q = 20.73, *p* < 0.0001), and striatum and Others (q = 8.060, *p* = 0.0007). \*\**p* < 0.01, \*\*\**p* < 0.001, \*\*\*\**p* < 0.0001. n = 3 mice. C, Representative images showing Rabies-GFP-labeled striatal input neurons, D1-tdTomato-labeled D1 medium spiny neurons, and merged fluorescence. Insets show enlarged examples of Rabies-GFP-positive and D1-tdTomato-positive neurons. Scale bars: 20 μm. D, Quantification of Rabies-GFP-labeled striatal input neurons classified as D1-positive or D1-negative. D1-MSNs constituted a significantly larger fraction of striatal inputs to mPFC-projecting NBM cholinergic neurons than D1-negative neurons. Paired t test, *t*_2_ = 4.356, *p* = 0.0489. n = 3 mice. Scale bars: 500 μm for low-magnification images and 10 μm for high-magnification images. **E,** Representative images showing Rabies-GFP-labeled input neurons in the ventral midbrain region together with tyrosine hydroxylase (TH) immunostaining and merged fluorescence. F, Quantification of Rabies-GFP-labeled neurons within the TH-positive dopaminergic region. Only a small fraction of Rabies-GFP-labeled input neurons overlapped with TH-positive neurons. Paired t test, *t*_10_ = 4.139, *p* = 0.0020, \*\**p* < 0.01. n = 11 slices containing VTA/SNc region from 3 mice.

Quantification of Rabies-GFP-labeled input neurons revealed a regionally biased input distribution (Figure 2B). Among the quantified regions, the striatum accounted for the largest proportion of monosynaptic inputs to mPFC-projecting NBM cholinergic neurons, followed by the cortex. Smaller fractions of input neurons were detected in the thalamus, hypothalamus, hippocampus, amygdala, and septum, with the remaining labeled neurons grouped as “Others.” These results indicate that mPFC-projecting NBM cholinergic neurons receive broad monosynaptic inputs, with a prominent contribution from the striatum.

Because the striatum represented the dominant input source, we next examined the identity of Rabies-GFP-labeled striatal neurons. Medium spiny neurons (MSNs) are the principal projection neurons of the striatum and are broadly divided into two major populations: D1 receptor-expressing MSNs, which are associated with direct-pathway “go” action, and D2 receptor-expressing MSNs, which are associated with indirect-pathway “no-go” action. These two MSN populations are present in approximately comparable proportions within the striatum. In ChAT-Cre;D1-tdTomato mice, tdTomato-positive neurons were used to identify D1-MSNs. Within the striatum, a subset of Rabies-GFP-positive neurons overlapped with D1-tdTomato fluorescence (Figure 2C), indicating that D1-MSNs provide monosynaptic input to mPFC-projecting NBM cholinergic neurons. Quantification showed that D1-positive neurons constituted a significantly larger fraction of Rabies-GFP-labeled striatal inputs than D1-negative neurons (Figure 2D), suggesting an enriched contribution from D1-tdTomato-positive striatal neurons within the labeled striatal input population (p < 0.05).

A substantial number of Rabies-GFP-labeled neurons was also observed in regions grouped as “Others,” including the ventral midbrain area containing the ventral tegmental area (VTA) and substantia nigra pars compacta (SNc). Because the VTA/SNc contains major dopaminergic neuronal populations, we next examined whether Rabies-GFP-labeled neurons in this region were dopaminergic. Immunostaining for tyrosine hydroxylase (TH), a marker of dopaminergic neurons, revealed only limited overlap between Rabies-GFP-labeled neurons and TH-positive cells (Figure 2E and F). These results indicate that although Rabies-GFP-labeled neurons were detected in the midbrain, TH-positive dopaminergic neurons represented only a small fraction of the labeled input neurons in this region.

Together, these results indicate that mPFC-projecting NBM cholinergic neurons receive widespread but regionally biased monosynaptic inputs, dominated by striatal inputs with an enriched contribution from D1-tdTomato-positive striatal neurons, whereas TH-positive dopaminergic neurons constitute only a minor labeled input population in the ventral midbrain.

### Rabies virus-mediated retrograde monosynaptic tracing of HDB cholinergic neurons projecting to the mPFC

We next applied the same projection-specific rabies tracing strategy to cholinergic neurons in the horizontal limb of the diagonal band of Broca (HDB), another major basal forebrain cholinergic region that innervates the mPFC. Similarly, a 1:1 mixture of AAV-DIO-TVA-mCherry and AAV-DIO-oG was injected into the HDB of ChAT-Cre;D1-tdTomato mice to express TVA-mCherry and oG in Cre-expressing cholinergic neurons (Figure 3A). Twenty-one days later, EnvA-ΔG-rabies-GFP (Rabies-GFP) was injected into the mPFC to label HDB cholinergic neurons projecting to the mPFC and their directly connected presynaptic inputs.

**Figure 3.**
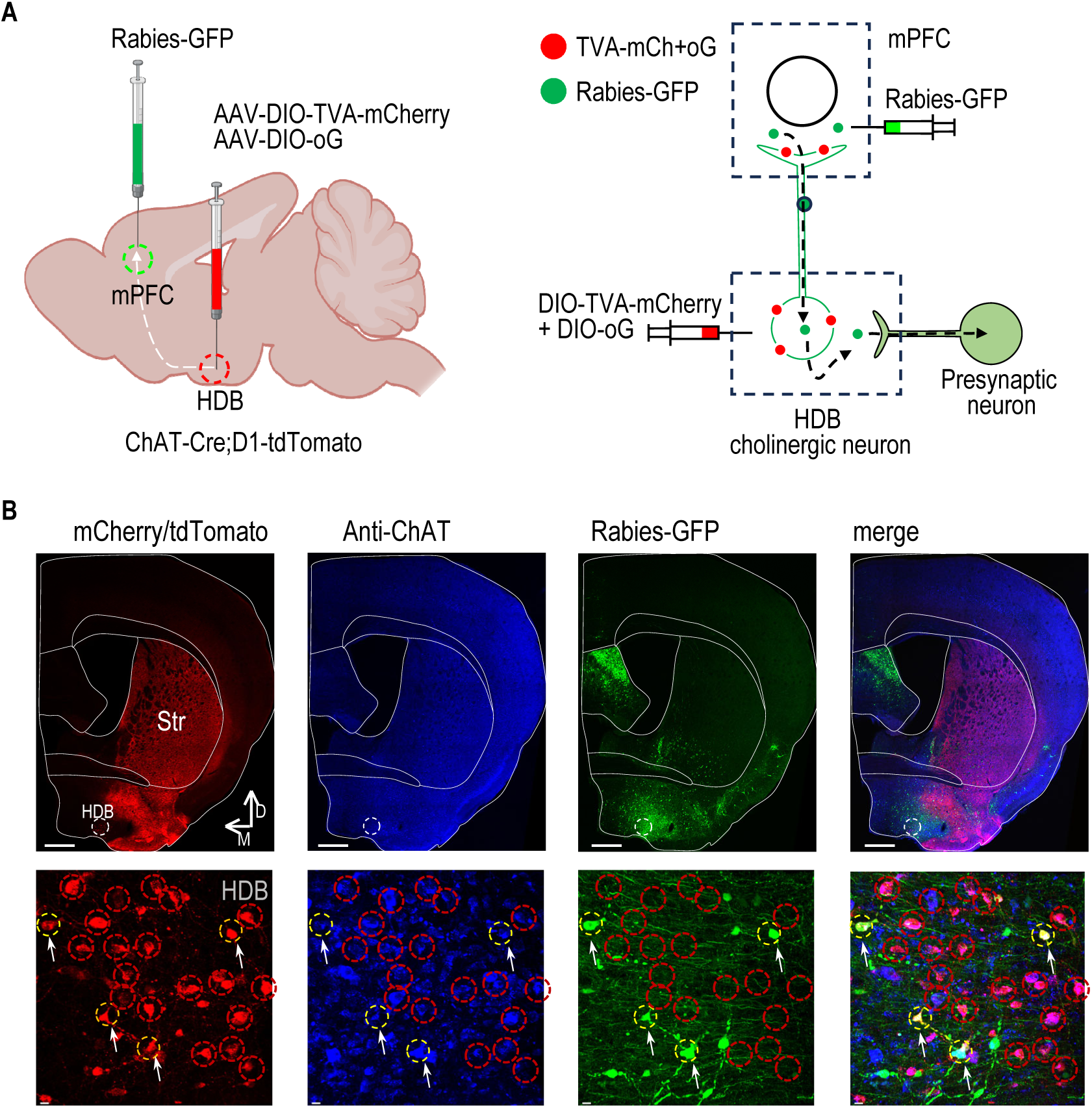
Rabies-mediated retrograde monosynaptic tracing of mPFC-projecting HDB cholinergic neurons. **A**, Schematic showing the viral injection strategy and tracing principle. Left, a 1:1 mixture of AAV-DIO-TVA-mCherry and AAV-DIO-oG was injected into the horizontal limb of the diagonal band of Broca (HDB) of ChAT-Cre;D1-tdTomato mice to selectively express TVA-mCherry and optimized rabies glycoprotein (oG) in cholinergic neurons. Twenty-one days later, EnvA-ΔG-rabies-GFP (Rabies-GFP) was injected into the medial prefrontal cortex (mPFC), the projection target of HDB cholinergic neurons. Right, Rabies-GFP selectively infected TVA-expressing axon terminals of HDB cholinergic neurons in the mPFC, was retrogradely transported to their somata in the HDB, and spread transsynaptically to directly connected presynaptic neurons in an oG-dependent manner. B, Representative images showing helper-virus expression, ChAT immunostaining, Rabies-GFP labeling, and merged fluorescence in the HDB. Low-magnification images show the injection region in the basal forebrain and Rabies-GFP-labeled cells. High-magnification images confirm the presence of starter cells. Red dashed circles indicate cholinergic neurons expressing TVA-mCherry/oG, and yellow dashed circles indicate starter cells co-expressing TVA-mCherry/oG and Rabies-GFP. Scale bars, 500 μm for low-magnification images and 10 μm for high-magnification images.

We also verified starter-cell labeling in the HDB using the same histological approach as in the NBM tracing experiment (Figure 3B). ChAT immunostaining revealed cholinergic neurons within the HDB, and a subset of ChAT-positive neurons overlapped with helper-virus-associated red fluorescence, confirming helper-virus targeting within the HDB cholinergic population. Among these ChAT-positive, red-labeled neurons, a subset was also positive for Rabies-GFP and was identified as starter cells. In addition, GFP-positive neurons lacking both ChAT immunostaining and red fluorescence were observed locally, consistent with local presynaptic neurons labeled by transsynaptic rabies spread from HDB cholinergic starter cells.

Together, these results confirmed successful projection-specific labeling of mPFC-projecting HDB cholinergic neurons, enabling comparison of whole-brain monosynaptic inputs to distinct basal forebrain cholinergic populations projecting to the same cortical target.

### Whole-brain mapping of inputs to HDB cholinergic neurons projecting to the mPFC

After confirming starter-cell labeling in the HDB, we again examined the whole-brain distribution of Rabies-GFP-labeled presynaptic neurons to identify direct inputs to mPFC-projecting HDB cholinergic neurons. Representative coronal sections arranged from rostral to caudal showed Rabies-GFP-labeled neurons distributed across multiple brain regions (Figure 4A). Labeled neurons were detected in the major regions quantified in this study, including the cortex, striatum, septum, thalamus, hypothalamus, hippocampus, and amygdala. Additional labeled neurons in regions not analyzed as separate categories were pooled into the “Others” group for quantification.

**Figure 4.**
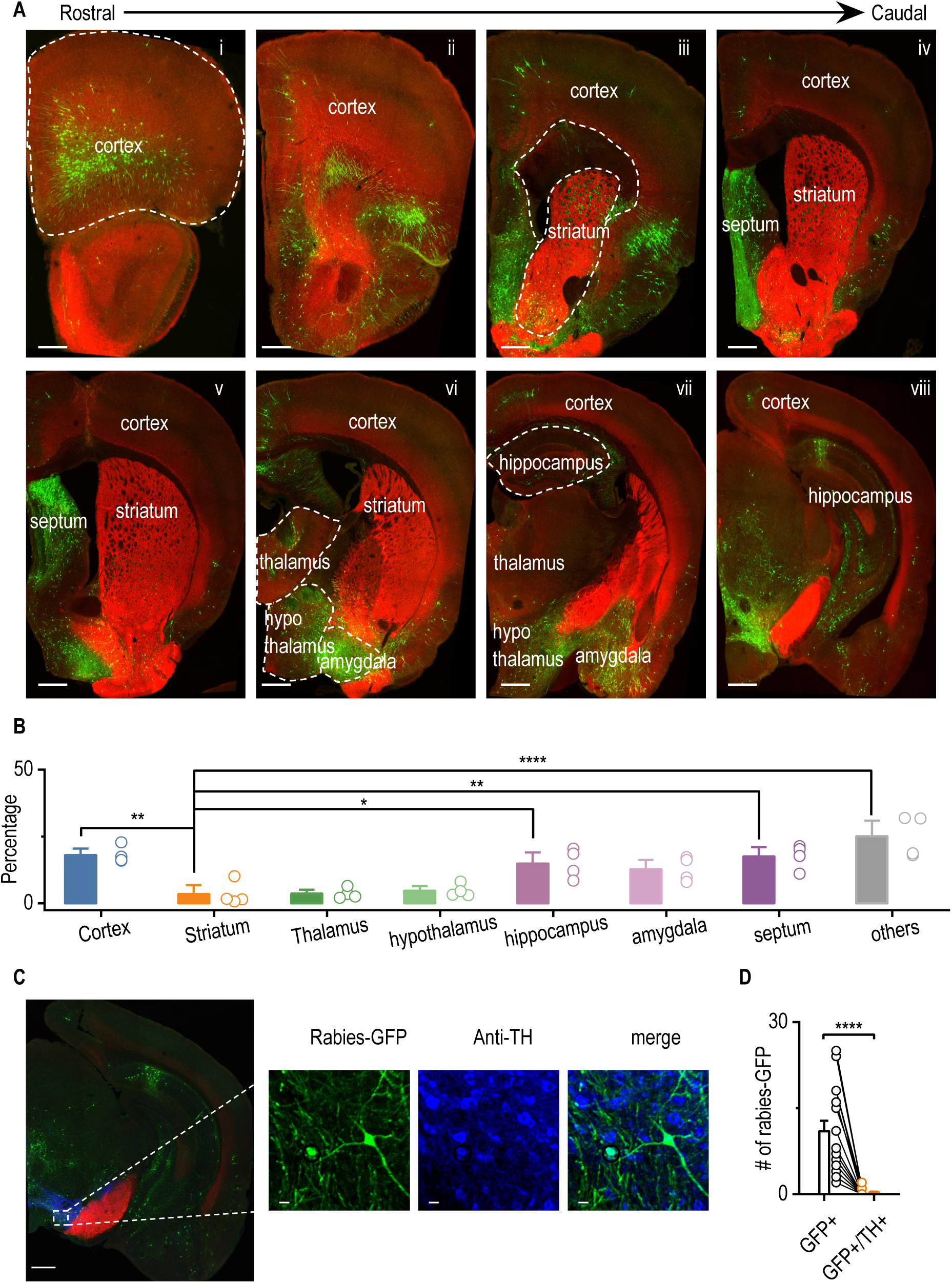
Whole-brain mapping of monosynaptic inputs to mPFC-projecting HDB cholinergic neurons. **A,** Representative coronal sections arranged from rostral to caudal showing Rabies-GFP-labeled monosynaptic input neurons throughout the brain. Cortex in sections i–viii, striatum in sections iii–vi, septum in sections iv–v, thalamus in sections vi–vii, hypothalamus in sections vi–vii, hippocampus in sections vii–viii, and amygdala in sections vi–vii. Additional labeled regions not analyzed as separate categories were pooled into the “Others” group for quantification. Dashed outlines indicate selected anatomical boundaries. Scale bars, 500 μm. B, Quantification of the relative distribution of Rabies-GFP-labeled input neurons across major brain regions. In contrast to the NBM tracing experiment, mPFC-projecting HDB cholinergic neurons received proportionally lower input from the striatum and relatively greater contributions from non-striatal regions, including cortex, septum, hippocampus, and amygdala. Regions not individually quantified were included in the “Others” category. One-way RM ANOVA, F_(7,21)_ = 10.15, *p* < 0.0001; Tukey’s post hoc comparisons, cortex and striatum (q = 6.260, *p* = 0.0038), striatum and hippocampus (q = 4.900, *p* = 0.0358), striatum and septum (q = 6.044, *p* = 0.0055), and striatum and Others (q = 9.268, *p* < 0.0001), \**p* < 0.05, ** *p* < 0.01, **** *p* < 0.0001. n = 4 mice. C, Representative images showing Rabies-GFP-labeled input neurons in the ventral midbrain region together with tyrosine hydroxylase (TH) immunostaining and merged fluorescence. Scale bars, 500 μm for low-magnification images and 10 μm for high-magnification images. **D,** Quantification of Rabies-GFP-labeled neurons within the TH-positive dopaminergic region. Only a small fraction of Rabies-GFP-labeled input neurons overlapped with TH-positive neurons. Paired t test, *t*_15_ = 6.028, *p* < 0.0001. \*\*\*\**p* < 0.0001. n = 16 slices containing VTA/SNc region from 4 mice.

Quantification of Rabies-GFP-labeled input neurons revealed that mPFC-projecting HDB cholinergic neurons received broadly distributed monosynaptic inputs across multiple brain regions (Figure 4B). In contrast to the NBM tracing experiment, striatal neurons contributed a relatively small fraction of the total labeled input population to HDB–mPFC cholinergic neurons. Instead, larger proportional contributions were observed from non-striatal regions, including the cortex, hippocampus, amygdala, septum, and regions grouped as “Others.” These results indicate that, although both NBM- and HDB-mPFC cholinergic neurons receive widespread monosynaptic inputs, the HDB-mPFC cholinergic population is less dominated by striatal input and instead receives a broader distribution of non-striatal afferents.

A subset of Rabies-GFP-labeled neurons was also observed in the ventral midbrain region containing the ventral tegmental area and the substantia nigra pars compacta. Because this region contains major dopaminergic neuronal populations, we examined whether the Rabies-GFP-labeled neurons were dopaminergic. Immunostaining for tyrosine hydroxylase (TH) revealed only limited overlap between Rabies-GFP-labeled neurons and TH-positive cells (Figure 4C,D), indicating that TH-positive dopaminergic neurons represented only a small fraction of the labeled input population in this region.

Together, these results demonstrate that mPFC-projecting HDB cholinergic neurons receive widespread monosynaptic inputs with an input distribution distinct from that of mPFC-projecting NBM cholinergic neurons, including proportionally weaker striatal input and broader contributions from non-striatal regions.

### Striatal inhibitory transmission onto NBM cholinergic neurons is reduced in 5xFAD mice

Identification of the striatum as the major monosynaptic input source to mPFC-projecting NBM cholinergic neurons suggested a prominent striatal–NBM pathway. We therefore next examined whether this anatomical connection forms functional synaptic input onto NBM cholinergic neurons and whether this functional connectivity is altered in the 5xFAD mouse model, given the vulnerability of NBM cholinergic neurons in Alzheimer’s disease (AD). To test this, we infused AAV-Chrimson into the striatum of ChAT-eGFP mice and performed whole-cell voltage-clamp recordings from eGFP-positive NBM cholinergic neurons in acute brain slices. Optical stimulation was then used to activate Chrimson-expressing striatal axons in the NBM.

Optical stimulation of striatal axons evoked postsynaptic transmission in NBM cholinergic neurons. Bath application of the GABA_A_ receptor antagonist picrotoxin (PTX, 100 μM) abolished optically evoked postsynaptic currents (Figure 5A–C), confirming that these responses were mediated by GABA_A_ receptors. We next assessed whether the striatal inhibitory responses were monosynaptic. Application of tetrodotoxin (TTX) strongly suppressed optically evoked inhibitory postsynaptic currents (oIPSCs), whereas subsequent application of 4-aminopyridine (4-AP) in the continued presence of TTX partially restored the responses (Figure 5D–F). This TTX-sensitive and 4-AP-rescuable response pattern supports the presence of monosynaptic striatal inhibitory input onto NBM cholinergic neurons.

**Figure 5.**
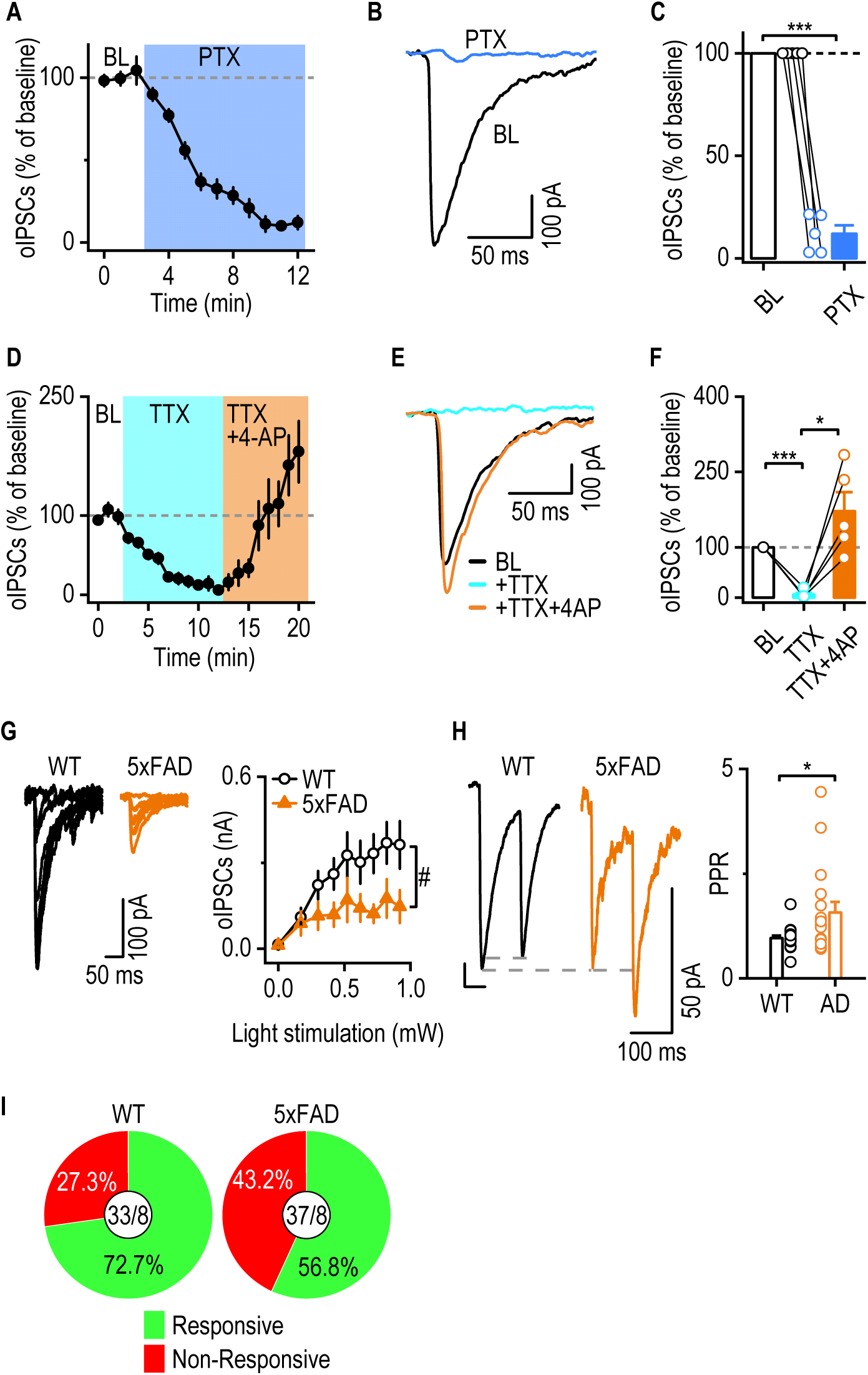
Striatal inhibitory transmission onto NBM cholinergic neurons is reduced in 5xFAD mice. **A,** Time course of optically evoked inhibitory postsynaptic currents (oIPSCs) recorded from ChAT-eGFP-positive NBM cholinergic neurons before and after bath application of picrotoxin (PTX, 100 μM). B, Representative oIPSC traces recorded at baseline and after PTX application. C, Quantification showing that PTX abolished oIPSCs. Paired t test, t_4_ = 21.38, *p* < 0.0001. \*\*\**p* < 0.001. n = 5 neurons from 4 mice. D, Time course of oIPSC amplitudes during baseline recording, tetrodotoxin (TTX) application, and subsequent TTX plus 4-aminopyridine (4-AP) application. E, Representative oIPSC traces recorded at baseline, in TTX, and in TTX plus 4-AP. F, Summary quantification showing that TTX strongly suppressed oIPSCs and that 4-AP restored light-evoked responses in the continued presence of TTX. Paired t test, t_4_ = 26.70, *p* < 0.0001 for BL vs TTX; t_4_ = 4.498, *p* = 0.0108 for TTX vs TTX+4AP. \**p* < 0.05, \*\*\**p* < 0.001. n = 5 neurons from 4 mice. G, Representative oIPSC traces showing striatal input to NBM cholinergic neurons. 5xFAD mice showed reduced oIPSC amplitudes compared with WT controls. Mixed-effects model, *F*_(1, 42)_ = 4.467, *p* = 0.0405. n = 23 neurons from 8 WT mice and 21 neurons from 8 5xFAD mice. ^#^*p* < 0.05. **H,** Representative traces of paired-pulse oIPSCs. Paired-pulse ratio of oIPSCs was evoked by two optical stimuli separated by a 100-ms inter-stimulus interval. 5xFAD mice showed an increased paired-pulse ratio compared with WT mice. Unpaired t test with Welch’s correction, t_17.67_ = 2.340, *p* = 0.0313. \**p* < 0.05. n = 20 neurons from 8 mice (WT) and 17 neurons from 8 mice (5xFAD). I, Pie charts showing the proportion of light-responsive and non-responsive NBM cholinergic neurons in WT and 5xFAD mice. WT: 72.7% responsive and 27.3% non-responsive, 33 neurons from 8 mice; 5xFAD: 43.2% responsive and 56.8% non-responsive, 37 neurons from 8 mice.

Having established functional inhibitory connectivity from the striatum to NBM cholinergic neurons, we next compared synaptic transmission between WT and 5xFAD mice. Input-output analysis showed that increasing opto-stimulation intensities evoked progressively larger oIPSCs in WT neurons, whereas oIPSC amplitudes were reduced in 5xFAD mice (Figure 5G). Paired-pulse analysis further revealed an increased paired-pulse ratio in 5xFAD mice compared with WT controls (Figure 5H), consistent with reduced presynaptic release probability at striatal inhibitory synapses onto NBM cholinergic neurons. Consistent with the reduction in oIPSC amplitude, we also observed a lower proportion of opto-responsive NBM cholinergic neurons in 5xFAD mice (Figure 5I).

Together, these findings demonstrate that striatal inputs form functional monosynaptic GABAergic synapses onto NBM cholinergic neurons and suggest that this pathway is impaired in 5xFAD mice, with reduced inhibitory synaptic strength, increased paired-pulse ratio, and a lower proportion of light-responsive NBM cholinergic neurons.

## DISCUSSION

In this study, we first used projection-specific rabies-mediated monosynaptic tracing to map upstream inputs to basal forebrain cholinergic neurons projecting to the mPFC. We found that mPFC-projecting NBM cholinergic neurons receive widespread but regionally biased monosynaptic inputs, with the striatum representing a major upstream source. Within this striatal input population, D1-expressing medium spiny neurons (D1-MSNs) contributed prominently to the labeled presynaptic neurons. In contrast, mPFC-projecting cholinergic neurons in the HDB showed a distinct input distribution, with proportionally weaker striatal input and broader contributions from non-striatal regions. We then used optogenetic electrophysiology to examine the functional striatal input to NBM cholinergic neurons. Optical activation of striatal terminals evoked monosynaptic GABAergic inhibitory postsynaptic currents in NBM cholinergic neurons, and this pathway was impaired in 5xFAD mice, as reflected by reduced inhibitory synaptic strength, increased paired-pulse ratio, and a lower proportion of light-responsive NBM cholinergic neurons.

The distinct input profiles observed across the NBM–mPFC and HDB–mPFC tracing experiments are consistent with previous evidence that basal forebrain cholinergic neurons are anatomically heterogeneous and organized into partially segregated circuit modules rather than a uniform diffuse projection system. Prior whole-brain tracing and input-output studies have shown that basal forebrain cholinergic circuits vary across cell types, subregions, and projection targets (Do et al., 2016). Our results further suggest that mPFC-projecting cholinergic neurons located in the NBM and HDB may receive different upstream influences. The prominent striatal input to NBM–mPFC cholinergic neurons indicates that this projection-defined population may be particularly positioned to integrate basal ganglia-related information. This finding is partly consistent with prior work showing the caudate-putamen as a major input source for several basal forebrain cholinergic output populations, although it differs from the relatively limited caudate-putamen input reported for broadly sampled mPFC-projecting basal forebrain cholinergic neurons (Do et al., 2016; Hu et al., 2016; Gielow and Zaborszky, 2017; Chen et al., 2025; Sun et al., 2026). This difference may reflect subregional specificity of the NBM cholinergic population, species differences, or differences in viral targeting and injection strategy.

The identity of midbrain inputs to basal forebrain cholinergic neurons is another important point raised by our study. We observed Rabies-GFP-labeled neurons in the VTA/SNc region, but only a small fraction of these cells overlapped with TH immunostaining. This finding is consistent with previous work showing that most ventral midbrain input cells to basal forebrain cholinergic neurons are TH-negative, despite the presence of occasional TH-positive input neurons (Gielow and Zaborszky, 2017). Because the VTA/SNc contains dopaminergic, GABAergic, and glutamatergic neuronal populations, our data suggest that the ventral midbrain input to mPFC-projecting basal forebrain cholinergic neurons is likely not predominantly dopaminergic (Van Bockstaele and Pickel, 1995; Yamaguchi et al., 2007). Future studies using cell-type-specific markers or intersectional viral approaches will be important for defining the neurochemical identity of these inputs and determining their functional contribution to basal forebrain cholinergic circuit regulation.

Our optogenetic electrophysiology experiments demonstrated that the anatomical striatal input identified by rabies tracing corresponds to a functional inhibitory pathway onto NBM cholinergic neurons. Consistent with the GABAergic identity of MSNs, optical activation of striatal terminals evoked GABA_A receptor-mediated oIPSCs in NBM cholinergic neurons, and the TTX/4-AP rescue experiment supported a monosynaptic component of this input (Gerfen and Surmeier, 2011; Day et al., 2024). These findings identify a functional striatal–NBM inhibitory pathway that may allow striatal activity to regulate basal forebrain cholinergic output to the mPFC. This pathway may be particularly relevant to AD, because degeneration or dysfunction of basal forebrain cholinergic neurons has long been associated with AD and cognitive decline (Whitehouse et al., 1982; Geula et al., 2021). In 5xFAD mice, striatal inhibitory transmission onto NBM cholinergic neurons was reduced, as reflected by smaller oIPSC amplitudes and increased paired-pulse ratio. Together with the lower proportion of light-responsive NBM cholinergic neurons, these results suggest that the striatal–NBM pathway is functionally weakened in 5xFAD mice, potentially involving reduced presynaptic release probability and/or loss of functional synaptic connectivity. This interpretation is consistent with recent work showing that AD mouse models exhibit long-range alterations in circuit connectivity beyond local pathology or cell loss (Ye et al., 2022; Ye et al., 2024; Ye et al., 2025).

An important question raised by these findings is how reduced striatal inhibitory transmission onto NBM cholinergic neurons can coexist with MSN hyperactivity observed in AD-related conditions (Huang et al., 2025; Huang et al., 2026). Increased somatic activity of MSNs may not necessarily predict stronger synaptic output at all downstream targets, because synaptic output also depends on presynaptic release probability, terminal plasticity, and synaptic integrity (Branco and Staras, 2009). Instead, AD-related circuit remodeling may be pathway- and neurotransmitter-specific. For example, previous work from our group showed enhanced mPFC-to-dorsomedial striatum (DMS) glutamatergic transmission in AD models, indicating that some long-range excitatory projections can be strengthened under AD-related conditions (Huang et al., 2025; Huang et al., 2026). In contrast, the present study shows weakened long-range GABAergic transmission from striatal MSNs to NBM cholinergic neurons. This contrast suggests that AD-related synaptic remodeling does not simply reflect a global increase or decrease in circuit activity, but may depend on the presynaptic cell type, neurotransmitter identity, projection target, and local synaptic environment (Barthet and Mulle, 2020; Meftah and Gan, 2023).

This pathway-specific interpretation may also help explain differences between striatal cholinergic interneurons and basal forebrain cholinergic projection neurons. In previous studies, striatal cholinergic interneurons showed robust inhibitory responses to MSN stimulation, and MSN-to-cholinergic interneuron (CIN) inhibition was enhanced in AD models, consistent with MSN hyperactivity (Gangal et al., 2023; Huang et al., 2026). In contrast, in the present study, only a subset of NBM cholinergic neurons responded to optogenetic activation of striatal terminals, and MSN-to-NBM inhibitory responses were weakened in 5xFAD mice. Thus, local MSN-to-CIN inhibition and long-range MSN-to-BF cholinergic projection neuron inhibition may be differentially affected in AD. Local MSN outputs may more directly reflect increased MSN excitability. In contrast, long-range striatal projections to the NBM may be more vulnerable to axonal dysfunction, reduced presynaptic release probability, or loss of functional synaptic contacts. Together, these findings suggest that AD-related circuit dysfunction occurs at multiple levels, including somatic excitability, neurotransmitter-specific synaptic remodeling, and pathway-specific long-range connectivity.

Although our study reveals broad monosynaptic input patterns to mPFC-projecting basal forebrain cholinergic neurons, more detailed subregional analyses would further improve understanding of input specificity. In the present study, inputs were quantified across major anatomical categories, including cortex, striatum, thalamus, hypothalamus, hippocampus, amygdala, and septum. Future studies subdividing these broad regions into defined subregions, cortical areas and layers, thalamic nuclei, or striatal territories would provide a more precise map of how distinct upstream networks converge onto NBM- and HDB-mPFC cholinergic circuits (Do et al., 2016; Hu et al., 2016; Gielow and Zaborszky, 2017; Lu et al., 2021; Chen et al., 2025). Moreover, previous rabies-tracing studies in AD mouse models have reported altered or reduced input connectivity in other brain regions, but whether similar anatomical connectivity changes occur in basal forebrain cholinergic projection neurons (BF-CPNs) remains unknown. Comparing rabies-labeled input organization between WT and 5xFAD mice would help determine whether the reduced striatal inhibitory transmission observed here reflects loss of presynaptic input neurons, altered input distribution, or synaptic dysfunction without major anatomical reorganization (Ye et al., 2022; Ye et al., 2024; Ye et al., 2025). It will also be important to determine whether additional factors, such as biological sex or sustained exposure to substances, further modify synaptic transmission or connectivity onto BF-CPNs. Moreover, because the present functional analysis was performed in the 5xFAD model, which primarily reflects Aβ-related pathology, it remains unclear whether similar striatal–NBM synaptic alterations occur in tau-based or mixed Aβ/tau pathological conditions. Future studies using additional AD models will be needed to determine whether weakening of striatal inhibitory input to NBM cholinergic neurons represents a shared feature of AD-related circuit dysfunction or an Aβ-associated pathway-specific alteration. Finally, the behavioral role of input–NBM/HDB–mPFC circuits remains to be established. Future pathway-specific manipulation during mPFC-dependent cognitive, motivational, or decision-making tasks will be critical for determining how these upstream inputs regulate basal forebrain cholinergic control of cortical function.

In summary, our study identifies projection-specific input organization of mPFC-projecting basal forebrain cholinergic neurons and reveals a prominent striatal input to the NBM–mPFC cholinergic pathway. By functionally validating this anatomical connection, we show that striatal inputs form monosynaptic GABAergic synapses onto NBM cholinergic neurons and that this inhibitory transmission is weakened in 5xFAD mice. These findings expand our understanding of basal forebrain cholinergic circuit organization and suggest that AD-related cholinergic dysfunction may involve disruption of upstream synaptic inputs in addition to intrinsic degeneration of cholinergic neurons.

## MATERIALS AND METHODS

### Animals

ChAT-eGFP mice (stock #007902), 5xFAD mice (stock #034848), ChAT-Cre mice (ChAT-IRES-Cre-δ-neo; stock #031661), and Drd1a-tdTomato mice (stock #016204) were obtained from The Jackson Laboratory. All mouse lines were maintained on a C57BL/6J background. ChAT-eGFP mice were crossed with 5xFAD mice to generate ChAT-eGFP;5xFAD mice for electrophysiological recordings from basal forebrain cholinergic neurons in an Alzheimer’s disease mouse model. ChAT-Cre mice were crossed with Drd1a-tdTomato mice to generate ChAT-Cre;Drd1a-tdTomato mice for projection-specific rabies tracing experiments. Mice were group-housed at 23°C under a 12-h light/dark cycle, with food and water available ad libitum. Both male and female mice were used in this study. All animal procedures were approved by the Texas A&M Institutional Animal Care and Use Committee and were performed in accordance with the National Research Council Guide for the Care and Use of Laboratory Animals.

### Confocal imaging and cell counting

Seven days after rabies virus injection, mice were intracardially perfused with 4% paraformaldehyde (PFA) in phosphate-buffered saline (PBS) to allow sufficient time for rabies virus replication and reporter expression. Brains were extracted, post-fixed overnight in 4% PFA/PBS, and dehydrated in 30% sucrose. Whole brains were serially sectioned into 50-μm coronal slices using a cryostat. Confocal images were acquired using a confocal laser-scanning microscope (Fluoview 3000, Olympus). Fluorescent image stacks were reconstructed in three dimensions, and cell counts were manually performed using Imaris 8.3.1, as previously reported (Wei et al., 2018). Labeled neurons were counted using the Spot module in Imaris, which was also used for colocalization analysis. Brain regions were identified and registered according to the Paxinos mouse brain atlas (Franklin and Paxinos, 2007).

### Quantification and normalization of Rabies-GFP⁺ neurons

Rabies virus–labeled (GFP⁺) neurons were manually counted across serial coronal sections obtained from each animal. Counting was performed separately for defined brain regions, including the cortex, striatum, septum, thalamus, hypothalamus, hippocampus, amygdala, ventral midbrain (VTA/SNc), and other labeled regions. To account for variability in regional coverage across serial sections, regional fractions were first calculated within each section and then averaged within each animal. Animals, not sections, were used as the biological replicates for group-level statistics.

For each section, the number of GFP⁺ neurons in each region was divided by the total number of GFP⁺ neurons detected in that section, yielding the fractional contribution of each region within that section. These per-section regional fractions were then averaged across all sections in which the region was present, resulting in a mean per-slice contribution for each region. Because not all regions were represented in every section, this procedure normalized for unequal sampling coverage across the series. The resulting mean regional fractions were further normalized so that the sum across all regions equaled 1, providing a relative measure of the distribution of rabies-labeled neurons independent of total labeling intensity. These normalized values were used to describe the overall distribution of input neurons across brain regions within each animal. When multiple animals were analyzed, normalized regional fractions were averaged across animals to obtain group means ± SEM for each region.

### Stereotaxic virus infusion

Stereotaxic viral infusions were performed as previously described (Wang et al., 2015; Huang et al., 2017; Ma et al., 2017; Roltsch Hellard et al., 2019; Cheng et al., 2021; Gangal et al., 2025; Xie et al., 2025; Xie et al., 2026). Mice were anesthetized and placed in a stereotaxic frame. The scalp was opened to expose the skull, and the bregma and lambda were identified. A three-axis micromanipulator was used to determine stereotaxic coordinates, and small craniotomies were made above the target injection sites according to the Paxinos mouse brain atlas. For projection-specific rabies tracing, ChAT-Cre;D1-tdTomato mice were used. rAAV8/CA-Flex-RG (UNC Vector Core, AV5005I) and rAAV8/EF1a-Flex-TVA-mCherry (UNC Vector Core, AV5008C) were mixed at a 1:1 ratio immediately before infusion. A total volume of 0.5 μL of the helper-virus mixture was bilaterally injected into either the nucleus basalis of Meynert (NBM; AP: −0.35 mm, ML: ±1.60 mm, DV: −4.60 mm) or the horizontal limb of the diagonal band of Broca (HDB; AP: +0.14 mm, ML: ±1.25 mm, DV: −5.80 mm). Twenty-one days after helper-virus injection, 0.5 μL of EnvA-pseudotyped, glycoprotein-deleted N2c rabies virus expressing GFP (EnvA-ΔG-N2c-Rabies-GFP; Thomas Jefferson University) was bilaterally injected into the medial prefrontal cortex (mPFC; AP: +1.94 mm, ML: ±0.35 mm, DV: −2.25 mm). For optogenetic electrophysiology experiments, rAAV8/syn-ChR-88m19-tdTomato (UNC Vector Core, AV5841D) was bilaterally injected into the striatum of ChAT-eGFP and 5xFAD;ChAT-eGFP mice (0.5 μL per side; AP: +0.50 mm, ML: ±1.70 mm, DV: −3.50 mm). All viral infusions were performed at a rate of 0.1 μL/min. To minimize backflow, microinjectors were left in place for 10 min after each infusion before being slowly withdrawn. The scalp was then sutured, and mice were allowed to recover before subsequent experiments.

### Electrophysiological recordings of brain slices

#### Brain slice preparation

Mice were deeply anesthetized, and brains were rapidly removed and placed in ice-cold cutting solution. Coronal basal forebrain slices containing the NBM were cut at 250 μm thickness. The cutting solution consisted of the following (in mM): 40 NaCl, 143.5 sucrose, 4 KCl, 1.25 NaH_2_PO4, 26 NaHCO_3_, 0.5 CaCl_2_, 7 MgCl2, 10 glucose, 1 sodium ascorbate, and 3 sodium pyruvate, and had a pH of 7.35 and an osmolarity of 305-310 mOsm. The solution was saturated with 95% O_2_ + 5% CO_2_. After sectioning, slices were incubated at 32°C for 45 min in a 1:1 mixture of cutting solution and external solution. The external solution, which had a pH of 7.35 and an osmolarity of 305-310 mOsm, was composed of the following (in mM): 125 NaCl, 4.5 KCl, 2 CaCl_2_, 1 MgCl_2_, 1.25 NaH_2_PO_4_, 25 NaHCO_3_, 15 sucrose, and 15 glucose. The external solution was continuously saturated with 95% O_2_ + 5% CO_2_. Slices were then maintained in external solution at room temperature until recording.

#### Electrophysiological recordings

For whole-cell recordings, basal forebrain slices were transferred to a recording chamber mounted on the fixed stage of an upright microscope (Olympus) and continuously perfused with oxygenated external solution at 32°C at a flow rate of approximately 2 mL/min. NBM cholinergic neurons were identified by eGFP fluorescence in ChAT-eGFP and 5xFAD;ChAT-eGFP mice and visualized using a 40× water-immersion objective and an infrared-sensitive CCD camera.

Whole-cell voltage-clamp recordings were performed using a Multiclamp 700B amplifier, Clampex 10.6 software, and a Digidata 1550A data acquisition system (Molecular Devices, Sunnyvale, CA). Recording pipettes were pulled from borosilicate glass capillaries (World Precision Instruments, Sarasota, FL) using a micropipette puller (Model P-97, Sutter Instrument) and had a resistance of 3–6 MΩ.

Optically evoked inhibitory postsynaptic currents (oIPSCs) were recorded from eGFP-positive NBM cholinergic neurons using a high-chloride intracellular solution. This solution contained the following components (in mM): 125 CsCl, 6 NaCl, 10 HEPES, 1 EGTA, 10 QX-314.Cl, 2 MgATP, 6 Na₃GTP, and 2 Na₂CrPO₄ with pH adjusted to 7.25 and osmolarity of 280 mOsm. Cells were voltage-clamped at −60 mV. Optical stimulation was delivered to activate striatal axon terminals expressing ChR-88m19-tdTomato in the NBM. To confirm that optically evoked currents were mediated by GABA_A_ receptors, oIPSCs were recorded before and after bath application of picrotoxin (PTX, 100 μM). To assess whether striatal inputs onto NBM cholinergic neurons contained a monosynaptic component, oIPSCs were recorded at baseline, after bath application of tetrodotoxin (TTX, 1μM), and after subsequent application of 4-aminopyridine (4-AP, 0.5 mM) in the continued presence of TTX. Suppression of oIPSCs by TTX and partial restoration by 4-AP were used to support monosynaptic optically evoked inhibitory transmission.

All patch-clamp recording data were analyzed using Clampfit (Molecular Devices).

### Immunohistochemistry

For ChAT immunostaining, free-floating brain sections were washed in 0.5% PBS-Triton X-100 (PBS-TX) and blocked with 5% bovine serum albumin (BSA) in 0.5% PBS-TX for 1 h at room temperature. Sections were incubated with goat anti-ChAT antibody (Millipore, AB144P; 1:100) diluted in 5% BSA/PBS-TX for two nights at 4°C. After three washes in 0.5% PBS-TX, sections were incubated with Alexa Fluor 647-conjugated donkey anti-goat IgG (H+L) secondary antibody (Invitrogen, A21447; 1:500) diluted in 5% BSA/PBS-TX for 2 h at room temperature. Sections were then washed three times in 0.5% PBS-TX before mounting and imaging.

For TH immunostaining, free-floating brain sections were washed in 0.5% PBS-TX and blocked with 5% BSA in 0.5% PBS-TX for 1 h at room temperature. Sections were incubated with chicken anti-tyrosine hydroxylase antibody (Abcam, ab76442; 1:1000) diluted in 5% BSA/PBS-TX overnight at 4°C. After three washes in 0.5% PBS-TX, sections were incubated with Alexa Fluor 647-conjugated donkey anti-chicken IgY (H+L) secondary antibody (Invitrogen, A78952; 1:500) diluted in 5% BSA/PBS-TX for 2 h at room temperature. Sections were then washed three times in 0.5% PBS-TX before mounting and imaging.

### Statistical analysis

Data were analyzed using paired or unpaired two-tailed t tests (with Welch’s correction when equal variance was not assumed), one-way ANOVA, one-way repeated-measures ANOVA, two-way repeated-measures ANOVA, or mixed-effects models, as appropriate. Post hoc comparisons were performed when applicable. Normality and equal variance assumptions were assessed before parametric analyses. Statistical significance was defined as p < 0.05. Statistical analyses were performed using GraphPad Prism, SigmaPlot, or SPSS. Data are presented as mean ± SEM.

## DECLARATION OF GENERATIVE AI USE

During the preparation of this work, the authors used ChatGPT to assist with polishing the manuscript’s language, correcting grammar, and correcting typos. AI generated no scientific results, data analysis, interpretation, or conclusions. After using this tool, the authors carefully reviewed and edited the content as needed and take full responsibility for the content of the published article.

## ACKNOWLEDGEMENTS

This research was supported by NIH grants U01AA025932, R01AA027768, and R01AA030293 to J.W.; the Texas A&M University Division of Research Targeted Proposal Teams (TPT) funding program to J.W.; and the TRSA John P. McGovern Fellowship 2024 to Y.H. We thank Thomas Jefferson University for providing the EnvA-N2c-rabies virus, which was supported by NIH grant P40OD010996. We appreciate Dr. Rahul Srinivasan and Roger Garcia for their help with TH staining.

## AUTHOR CONTRIBUTIONS STATEMENT

J.W. and Y.H. conceived the project and designed the experiments. Y.H. and X.X. performed the stereotaxic surgery. Y.H. performed the electrophysiological experiments and analyzed the corresponding data. Y.H., M.F. and Z.L. performed the histology experiments. X.W. performed animal breeding for all experiments. J.W. and Y.H. wrote the manuscript.

